# Intracellular force comparison of pathogenic KIF1A, KIF5, and dynein by fluctuation analysis

**DOI:** 10.1101/2021.09.12.459977

**Authors:** Kumiko Hayashi, Shiori Matsumoto, Takuma Naoi, Yuki Idobata

## Abstract

In mammalian cells, there exist approximately 40 types of microtubule motor proteins that are assigned to specific cargo deliveries. For example, the kinesin-1 family motor KIF5 is the major motor responsible for anterograde mitochondrial transport, whereas the kinesin-3 family motor KIF1A is responsible for synaptic vesicle precursor transport. In contrast, cytoplasmic dynein is responsible for retrograde transport of nearly all cargos. The force and velocity of these microtubule motors have been investigated in *in-vitro* single-molecule experiments. In the present study, we compared the intracellular force and velocity of various types of motors in the mammalian neuronal axon obtained by non-invasive force measurement (fluctuation analysis) and extreme value analysis with those obtained by previous single-molecule experiments. As we found a high correlation between our results and the previous results, we next investigated synaptic vesicle precursor transport by hereditary spastic paraplegia-associated KIF1A variants (V8M, R350G, and A255V). KIF1A-V8M and KIF1A-A255V exhibited force and velocity impairment in mammalian neuronal axons, whereas the physical property of KIF1A-R350G was similar to that of the wild type. We believe that the development of new analytical techniques for investigating intracellular cargo transports is helpful to elucidate the molecular mechanism of KIF1A-associated neurological disorders.

**Statement of Significance:** Recent *in-vitro* single-molecule experiments have clearly revealed that microtubule motors only fully exert their functions when fully equipped with the proteins associated with cargo vesicle transport. This emphasizes the significance of intracellular physical measurements, in which the motors should fully function. In addition to motor force and velocity, the number of motors transporting a single cargo together is an important physical quantity to characterize cargo transport, but is difficult to estimate using *in-vitro* single-molecule experiments. In this study, we aimed to extract physical information on microtubule motors in the intracellular environment.

## Introduction

A large variety of cargo vesicles and organelles, such as endosomes, mitochondria, and phagosomes, are transported by microtubule motor proteins, kinesin and dynein, along microtubules spread throughout the cell, to deliver materials synthesized near the center of the cell to areas where they are required. In mammalian cells, there exist approximately 40 types of microtubule motor proteins that are assigned to specific cargo deliveries. For example, the kinesin-1 family motor KIF5 is the major motor responsible for anterograde transport of mitochondria (1), whereas the kinesin-3 family motor KIF1A is responsible for synaptic vesicle precursor (SVP) transport (2). In contrast, cytoplasmic dynein is responsible for retrograde transport of nearly all cargos (3).

The physical properties of the various microtubule motors, including their force and velocity, have been investigated in detail in *in-vitro* single-molecule experiments, typically using total internal reflection fluorescence (TIRF) microscopy (4–6) and optical tweezers (7–9). Recent studies have reported that forces differ among motors and that force and velocity are affected by pathogenic variants or the presence/absence of adaptor proteins associated with motor-cargo binding (10–14). The significant findings obtained by using *in-vitro* single-molecule techniques emphasize the significance of intracellular physical measurements, as microtubule motors were found to only fully exert their functions when fully equipped with their cargo vesicle transport-associated proteins, which is indeed realized in cells. Therefore, we aimed to investigate whether it would be possible to measure physical parameters of the motors directly in intracellular environment.

The time course of a specific cargo’s position in a cell can be followed in a low-invasive manner by fluorescence microscopy through attaching fluorescent labels to the cargo. However, unlike in *in-vitro* single-molecule experiments, it is difficult to extract physical information on motors from *in-vivo* time-course data because of the different cargo shapes and complex non-equilibrium environment. In recent years, to overcome these problems in time-course analysis under non-equilibrium conditions, we have developed a method for non-invasive force measurement of intracellular cargo transport based on time-course analysis using the statistical property of cargo fluctuation, obtaining idea from the non-equilibrium statistical mechanics (15–20).

As single-molecule experiments (10, 11, 14) have revealed force differences among KIF1A, KIF5, and dynein, we investigated anterograde SVP transport by KIF1A, mitochondrial transport mainly by KIF5, and retrograde transport by dynein in the axons of mammalian neurons. The validity of the force index of the non-invasive force measurement was confirmed by the high correlation between values and motor force values obtained by *in-vitro* single-molecule experiments. Encouraged by this finding, we aimed to investigate the pathogenic mutant KIF1A-V8M, which causes hereditary spastic paraplegia (21), known as a KIF1A-associated neurological disorder (KAND). Force generation impairment of this variant has been recently reported based on a single-molecule experiment; the V8M mutation is located in the motor domain of KIF1A, and molecular dynamics simulation predicted that the mutation alters neck-linker docking and catalytic-site closure, thus impairing force generation (11). Other hereditary spastic paraplegia-associated KIF1A variants (R350G and A255V) (22), whose phenotypes are similar to that of V8M (23), were also investigated using the force index *χ*.

As the KIF1A-V8M variant also leads to velocity impairment (11), we investigated the velocity of moving cargos using extreme value analysis (EVA) (24). Even in a highly viscous cellular environment, we could obtain a velocity value close to that under the no-load condition by using EVA, which has been applied to natural disasters (25), finance (26), sports (27), and human lifespan (28). The extreme velocity value obtained by EVA represents a significant quantitative measure for characterizing the true features of a variant under the no-load condition, as the velocity is expected to be readily decreased in the presence of cargos in the highly viscous cellular environment, especially when motor force generation is impaired.

*In-vitro* single-molecule experiments using TIRF microscopy and optical tweezers have provided substantial information on the force and velocity of microtubule motors. However, we believe that it is also important to develop new analytical techniques for obtaining time-course data of intracellular cargo transport to extract hidden information on their physical parameters. The measurement of intracellular physical parameters would be helpful to quantitatively understand the cellular phenomena associated with impaired cargo transport, *e.g*., abnormality in synaptogenesis due to impaired SVP transport because of KIF1A/UNC-104 (a homolog of KIF1A) mutation (12, 29). Studying microtubule motors from both aspects, *i.e*., *in-vitro* single-molecule experiments and intracellular time-course analyses, would promote the elucidation of intracellular cargo transport mechanisms.

## Materials and Methods

### Primary culture and transfection of neurons

Primary culture of hippocampal neurons was performed as described previously (30, 31), with some modifications. Although hereditary spastic paraplegia is a motor neuron disease, we used hippocampal neurons as they are easier to culture than motor neurons. Hippocampi of wild-type C57BL/6 mice (Japan SLC, Hamamatsu, Japan) at embryonic day 17 were dissected, and neurons were cultured in glass-bottom dishes (MatTek, Ashland, MA, USA) in culture medium (NbActiv4; BrainBits, Springfield, MA, USA), as described previously (30). After 4–7 day culture, the neurons were transfected with the plasmid vector for green fluorescence protein (GFP) fused human KIF1A (WT, R350G, V8M, A255V), provided by Dr. Shinsuke Niwa (Frontier Research Institute for Interdisciplinary Sciences, Tohoku University) (23), to observe SVP transport and the pDsRed2-Mito vector (BD Bioscience Clontech, Oxford, UK) to observe mitochondrial transport, using the calcium phosphate method (Takara Bio, Shiga, Japan). All animal experiments complied with the protocols approved by the Institutional Animal Care and Use Committee of Tohoku University (2016EgA-003, 2019EgA-001).

### Fluorescence microscopy

Cargo movement was observed under a fluorescence microscope (IX83; Olympus, Tokyo, Japan) equipped with a heating plate (CU-201; Live Cell Instrument, Seoul, Korea) maintained at 37°C. High power LED source (UHP-Mic-LED470; Prizmatix Ltd., Israel) was used for the observation. For fluorescence observation, B-27 supplement (Thermo Fisher Scientific, Chelmsford, MA, USA) and 150 μM 2-mercaptoethanol (Wako Chemical, Osaka, Japan) were added to the medium. Images were acquired using a 150× objective lens (UApoN 150×/1.45; Olympus, Tokyo, Japan) for SVP transport and a 100× objective lens (UPlanFLN 100×/1.30; Olympus) for mitochondrial transport, and an sCMOS camera (OLCA-Flash4.0 v.2.0; Hamamatsu Photonics, Hamamatsu, Japan) at 100 frames/s. In our previous study (18), we investigated data obtained at recording rates of 100 and 200 frames/s, and 100 frames/s was found to be sufficient for the following fluctuation analysis. Data for kymographs were obtained at 30 frames/s. The center positions of individual cargo vesicles were determined from the images using a custom software written in LabVIEW 2015 (National Instruments, Austin, TX, USA), as described previously (15). The kymographs in Fig. S3E in the Supplemental Information were obtained using ImageJ software (National Institutes of Health, Bethesda, MD, USA)(32).

The primary neuron cultures were repeated 27 times in the present study. Data for fluctuation analyses were collected from 11–14 preparations and from 38–50 cells for each type of motor. Data for EVA were collected from 5–6 preparations and from 58–64 cells for each type of motor. The cells used for observation were randomly chosen after visual inspection.

The precision of the positional measurements was verified using 300-nm latex beads (PolyScience, Niles, IL, USA) that were similar in size to the cargos. The standard deviation of a bead tightly attached to the glass surface was 8.3 ± 1.2 nm (*n* = 4 beads).

### Fluctuation analysis based on calculation of *χ*

Figure 1A shows a schematic diagram of SVP transport by GFP-labeled human KIF1A. As illustrated, a single SVP cargo is transported by multiple motors that exert a transport force *F*, which was our target of measurement. Figure 1B shows a typical constant velocity segment (CVS) in the time course of a cargo’s position. The anterograde direction was set as the positive direction of the *x*-axis. CVSs in time courses with velocities greater than 100 nm/s and lasting for more than 0.5 s were typically used for the following analyses.

**FIGURE 1.**
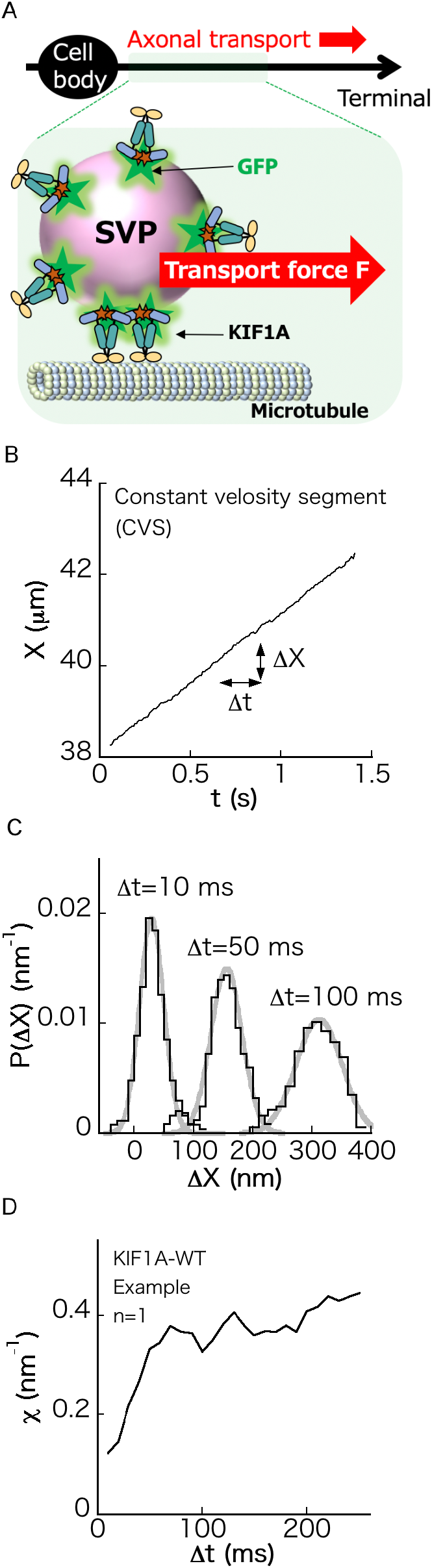
Synaptic cargo transport in the axon of a neuron. (A) A synaptic vesicle precursor (SVP) cargo (pink circle) is anterogradely transported by KIF1A motors labelled with GFP (green stars) along microtubules (green cylinder). The transport force (*F*) is generated by the motors. (B) Time course of the center position *X* of a moving SVP cargo in a constant velocity segment (CVS). The anterograde direction was set as the positive direction of the *x*-axis. Δ*X* = *X*(*t* + *Δt*) − *X*(*t*), where Δ*t* is a time interval. Δ*X* is a fluctuating quantity. (C) Distribution *P*(Δ*X*) of Δ*X* for Δ*t* = 10 ms, 50 ms, and 100 ms. *P*(Δ*X*) showed a diffusional behavior. *P*(Δ*X*) fit a Gaussian distribution (gray curve). (D) Example of *χ* (*n* = 1) (Eq. 3 in the Materials and Methods) calculated for the CVS (Fig. 1B) plotted as a function of Δ*t*.

Δ*X* (Fig. 1B) is a fluctuating quantity because the cargo’s position fluctuates due to thermal noise and collisions with other vesicles and cytoskeletons in the intracellular environment. CVS fluctuation Δ*X* in one direction was considered to be minimally affected by motors moving in the opposite direction, unlike in a tug-of-war segment (33), based on our previous observations using the dynein inhibitor, dynarrestin (18). Further, the activity of opposing motors at CVSs may be suppressed by adaptor proteins of cargo transport to avoid futile tug-of-war (34).

The distribution *P*(Δ*X*), where Δ*X* = *X*(*t* + Δ*t*) − *X*(*t*) (Fig. 1B) is plotted for Δ*t* = 10, 50, and 100 ms (Fig. 1C). Note that CVSs were chosen so that Δ*X* calculated from them belonged to the same statistical population (see Supplemental Information in (18)).

The force index *χ* introduced in our previous studies (15–18) was originally defined based on the idea of the fluctuation theorem (19, 35–37), as follows:

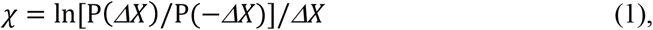

from the distribution *P*(Δ*X*) of displacement Δ*X* = *X*(*t* + Δ*t*) − *X*(*t*) (Fig. 1B).

*P*(Δ*X*) was fitted to a Gaussian function (Fig. 1C).

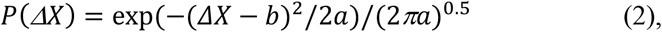

where the fitting parameters *a* and *b* correspond to the variance and mean of the distribution, respectively. By inserting Eq. 2 into Eq. 1, we obtain:

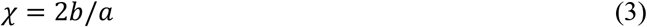

Equation 3 is equivalent to the equation *χ* = *v*/*D* when the diffusion coefficient *D* is defined as *D* = *a*/2Δ*t*.

A smoothing filter was applied to the values of *χ* to reduce the variation in the raw data (noisy data) for *χ* as a function of Δ*t*. We used the following simple averaging filter:

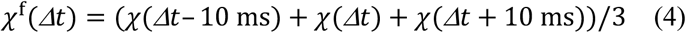

Note that *χ*^f^(Δ*t*) = (*χ*(Δ*t*) + *χ*(Δ*t* + 10 ms))/2 for Δ*t* = 10 ms and *χ*^f^(Δ*t*) = (*χ*(Δ*t* − 10 ms) + *χ*(Δ*t*))/2 for Δ*t* = 250 ms. The software used to calculate *χ* is available from a website (38). The behavior of *χ* as a function of Δ*t*, shown in Fig. 1D, converging to a constant value as Δ*t* → ∞, was explained in detail in our previous papers (15–20).

### Classification of *χ*-Δ*t* plots

Affinity propagation (39, 40), an exemplar-based clustering method, was adopted to cluster the two-dimensional data *χ*^f^(Δ*t* = 230 ms), *χ*^f^(Δ*t* = 250 ms or 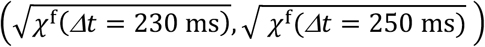. The square root of *χ* is often used to apply the cluster method to make the distance between adjoining clusters constant, as the value of *χ* has been observed to sometimes exponentially increase as the number of force-producing units (FPUs) increases, although the reason for this is unclear. The clustering method was applied using the ‘APCluster’ package in R (41). Clustering was stable for a wide range of values for the sole parameter (*q*) of affinity propagation analysis.

Because the result of clustering depended on the sample size (*n*) of *χ*, the stability of the cluster number against sample size was checked in a bootstrapping manner by performing cluster analysis 20 times for 75% of the original samples chosen randomly from the original samples. The probabilities of the numbers of clusters obtained in the 20 bootstrapping runs are shown in Figs. 2C, 4C, and 5C.

**FIGURE 2.**
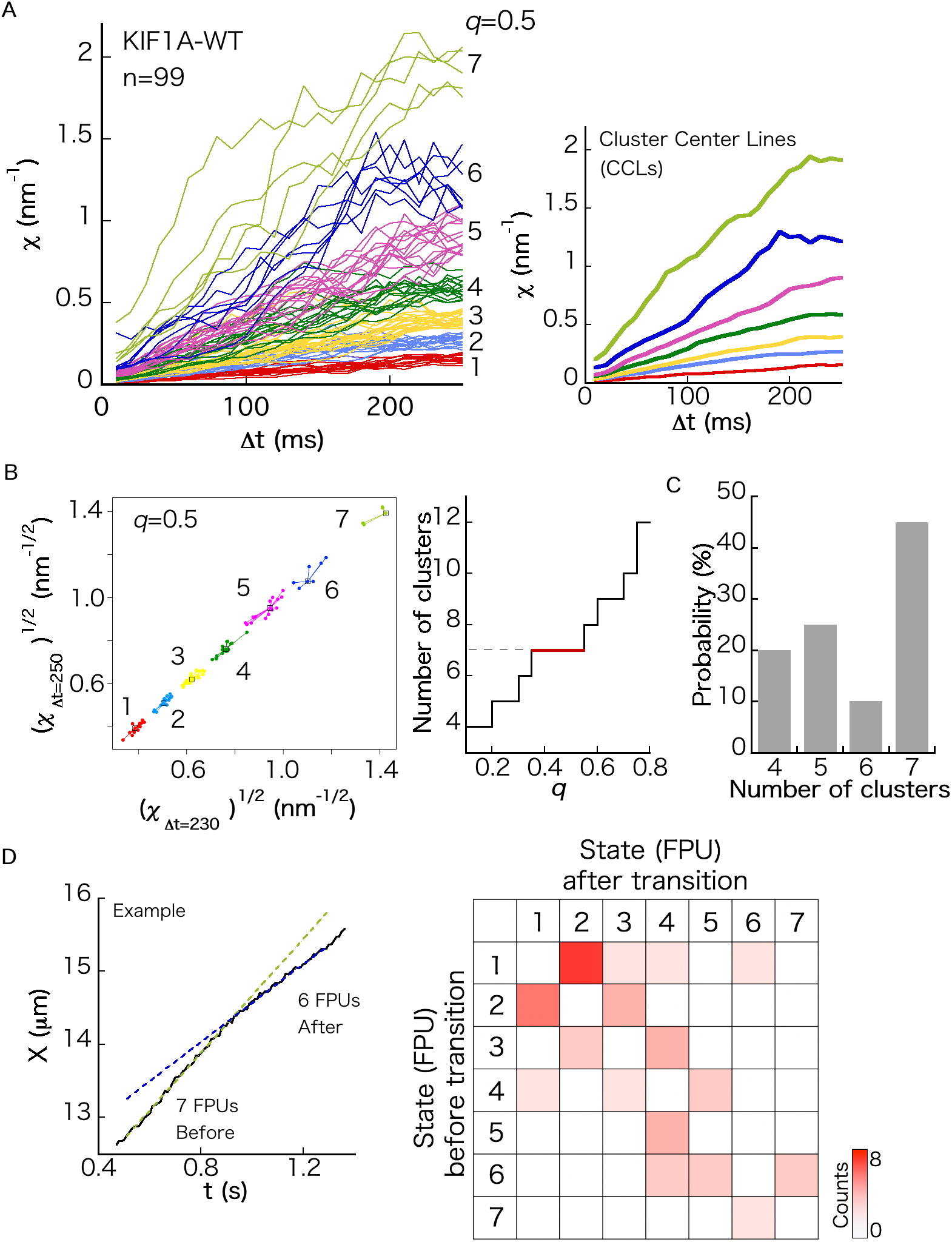
Force index for SVP transport by KIF1A-WT. (A) Left panel, *χ* as a function of Δ*t* for *n* moving cargos (*n* = 99). Each color represents the cluster assigned to each FPU. Right panel, cluster center lines (CCLs) calculated as the mean value of each cluster. (B) Result of affinity propagation analysis (see Materials and Methods). Cluster numbers are plotted as a function of *q*, which is the parameter of the cluster analysis. The most probable cluster number was determined as the cluster number valid for a wide range of *q*. (C) Result of boot-strapping analysis (see Materials and Methods). (D) Left panel, example time course of the center position *X* of a moving SVP cargo, containing a velocity change of adjacent CVSs. The straight lines are fitting lines. The number of each FPU at each CVS was determined by calculating *χ*. Right panel, heatmap of transition probability between states (*i.e.*, numbers of FPUs transporting a cargo) of adjacent CVSs.

### Statistical analysis

Welch’s *t*-test was applied to the data in Fig. 6A using the function (t.test()) in the R software (41). \**p* < 0.05, \*\**p* < 0.01, and \*\*\**p* < 0.001.

### Extreme value statistics (EVA)

First, the velocity values of moving cargos were collected from kymographs (*n* = 229 for KIF1A-WT, *n* = 334 for KIF1A-R350G, *n* = 190 for KIF1A-V8M, and *n* = 229 for KIF1A-A255V), and the data were divided into tens. Then, the largest value (*v*_max_) of the ten values was chosen. The probability distributions of *v*_max_ (*g*(*v*_max_)) for KIF1A-WT, KIF1A-R350G, KIF1A-V8M, and KIF1A-A255V are shown in Fig. S7 in the Supplemental Information. *g*(*v*_max_) = *dGEV*(*v*_max_)/*dv*_max_, where *GEV* is called the generalized extreme value distribution (a Weibull distribution) (24) and is written as:

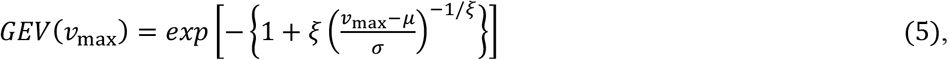

where *ξ*, *μ*, and *σ* are the parameters of *GEV*(*v*_max_). *GEV*(*v*_max_) obtained from the experimental data was fitted using Eq. 5 using the ‘ismev’ and ‘evd’ package sin R (41). By checking the convexity of the corresponding return level plot for the data, *GEV*(*v*_max_) was classified as a Weibull distribution. The extreme velocity value was then estimated as

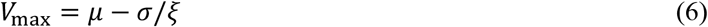

Note that the return level plot is defined as 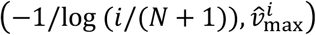 where 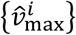 is the rearranged data of 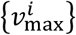 so that 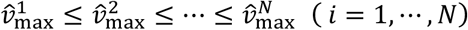, where *N* = *n*/10.

## Results and Discussion

### Force index for SVP transport by KIF1A-WT

We plotted the force index *χ* values (Eq. 3 in the Materials and Methods) for 99 anterograde moving cargos labeled with GFP-fused human KIF1A-WT (Fig. 2A, left panel). After applying the classification methods (Fig. 2B and 2C), we identified seven clusters of the force index *χ*, as shown in Fig. 2A, which were considered to correspond to seven force producing units (FPUs) consisting of KIF1A motors (likely, seven dimers), transporting a single SVP cargo. Here, the term “FPUs” is used rather than “motors” because the components of 1 FPU have not been clarified yet, although 1 FPU is considered to be a kinesin dimer for anterograde transport. Notably, the number of FPUs (7 FPUs) in the present study was larger than that previously reported for synaptic cargos labeled with GFP-fused synaptotagmin (6 FPUs) (18). Because GFP-fused human KIF1A-WT was expressed in the present study, there may have been excess KIF1A in a cell, leading to an increase in the number of FPUs. Apart from this difference in the number of FPUs, each value of *χ* for each FPU matched our previously reported results (18), demonstrating that there was no significant difference in *χ* between human KIF1A-WT and KIF1A originally derived from mice. In the right panel in Fig. 2A, the cluster center line, calculated as the mean value of each cluster of *χ*, was plotted for convenience in comparing the values with those of other motors.

To confirm the validity of the seven clusters identified by the classification methods (Fig. 2B and 2C), we investigated the cargo velocities of adjacent CVSs (Fig. 2D, left panel). We assumed that a velocity change of a moving cargo was followed by a change in the number of FPUs transporting the cargo (Fig. S1A in Supplemental Information). This assumption has been previously adopted by other groups (42, 43). For example, the velocity change in Fig. 2D was interpreted as a state transition of a cargo being transported by 7 FPUs to being transported by 6 FPUs, where the 7 FPUs and 6 FPUs were assigned by calculating *χ*. By repeating this procedure for many SVP cargos that contained two adjacent CVSs, the transition probabilities between the states of a moving SVP were measured (Fig. 2D, right panel). We found that there were no transitions within the same FPU group (cluster), and transitions mainly occurred between the adjacent numbers of FPUs. As it is natural for the number of FPUs to change by ±1, the classified clusters in Fig. 2A, left panel, seem to correctly describe the number of FPUs transporting an SVP cargo.

### Rescaled velocities for SVP transport

Previous studies focused on velocity changes observed in time courses of intracellular cargo transport, counting the number of motors carrying an intracellular cargo vesicle (42). The basic idea of this approach is depicted in Fig. S1A in the Supplemental Information. According to this idea, the velocity of a cargo changes when the number of motors (FPUs) changes as the velocity obeys Stokes’ law *F* = *Γv*, where *F* is the transport force exerted by the motors, *v* is the velocity of a cargo, and Γ is the friction coefficient of a cargo (Fig. S1A). In the previous study (42), the authors observed a sufficiently long trajectory of a moving cargo to determine the minimum velocity, and divided the other velocities in the time course by the minimum velocity to reduce the dependence of cargo size on velocity (note that cargo sizes vary, as shown in Fig. S1B in the Supplemental Information). They found several peaks in the histogram of the rescaled velocity, which were thought to correspond to the number of motors carrying the cargo (42).

Similarly, in the present study, we first identified the CVS assigned to 1 FPU by calculating *χ* and setting the velocity of the segment as *v*_1_, and then rescaled other velocities in CVSs of the time course as *v*/*v*_1_ (Fig. 3A). Although *v*_1_ often corresponded with the minimum velocity in a time course, many time courses did not comprise *v*_1_. Thus, *v*_1_ assigned by *χ* is more appropriate for rescaling other velocities.

**FIGURE 3.**
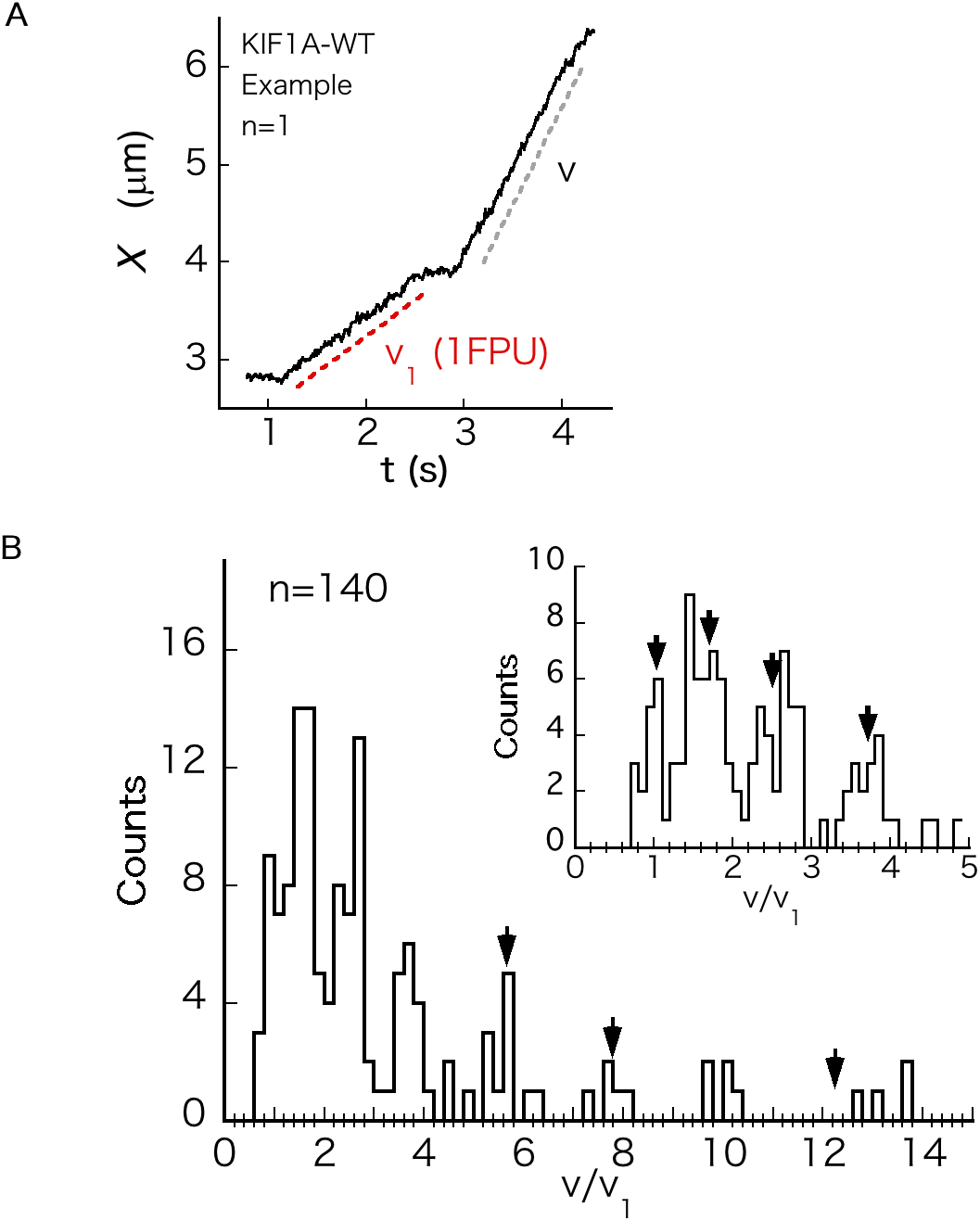
Analysis of velocity changes. (A) Example time course of the center position *X* of a moving SVP cargo, containing a CVS at velocity *v*_1_ of SVP transport by 1 FPU. *v*_1_ was found by calculating *χ*. *v* is the velocity at another CVS in the time course. Histogram of *v*/*v*_1_. The inset is a magnified view of the histogram for *v*/*v*_1_ < 5. The black arrows represent 〈*χ*_1_〉/〈*χ*_1_〉 (= 1), 〈*χ*_2_〉/〈*χ*_1_〉, 〈*χ*_3_〉/〈*χ*_1_〉, 〈*χ*_4_〉/〈*χ*_1_〉, 〈*χ*_5_〉/〈*χ*_1_〉, 〈*χ*_6_〉/〈*χ*_1_〉 and 〈*χ*_7_〉/〈*χ*_1_〉, where 〈*·*〉 represents the mean of *χ*_i_ belonging to the *i*-th cluster for Δ*t* = 250 ms.

After investigating long time courses of many moving cargos, we plotted a histogram of the rescaled velocities, *v*/*v*_1_, and found peaks (Fig. 3B). The inset in Fig. 3B shows a magnified view of the histogram for *v*/*v*_1_ < 5. The black arrows in Fig. 3B correspond to 〈*χ*_1_〉/〈*χ*_1_〉 (= 1), 〈*χ*_2_〉/〈*χ*_1_〉, 〈*χ*_3_〉/〈*χ*_1_〉, 〈*χ*_4_〉/〈*χ*_1_〉, 〈*χ*_5_〉/〈*χ*_1_〉, 〈*χ*_6_〉/〈*χ*_1_〉, and 〈*χ*_7_〉/〈*χ*_1_〉, where 〈·〉 represents the mean of *χ*_i_ belonging to the *i*-th cluster for Δ*t* = 250 ms (*i* = 1, …,7). Considering that *χ* = *v*/*D* (Eq. 3 in the Materials and Methods), where *D* is the diffusion coefficient of a moving cargo, *χ* is considered another rescaled velocity, and peaks of 〈*χ*_*i*_〉/〈*χ*_1_〉 correspond to the peaks in the histogram of *v*/*v*_1_. In conclusion, conceptually, our idea of determining the number of FPUs for intracellular cargo transport by using the force index *χ* is the same as using the velocity changes of cargos to this end, as previously suggested (42).

### Force index for SVP transport by KIF1A-V8M

Next, we investigated SVP transport under the condition that human KIF1A-V8M was expressed in the neurons. Although KIF1A originally derived from mice was mixed with KIF1A-V8M, the *χ* values for KIF1A-WT were largely different (Fig. 4 A, left panel). The thick black lines in Fig. 4A (right panel) represent cluster center lines (CCLs) for KIF1A-WT (Fig. 2A, right panel). The values of *χ* for KIF1A-V8M were approximately half of those for KIF1A-WT. As expected from the force impairment of KIF1A-V8M reported in an *in-vitro* single-molecule study (11), the values of *χ* were significantly decreased.

**FIGURE 4.**
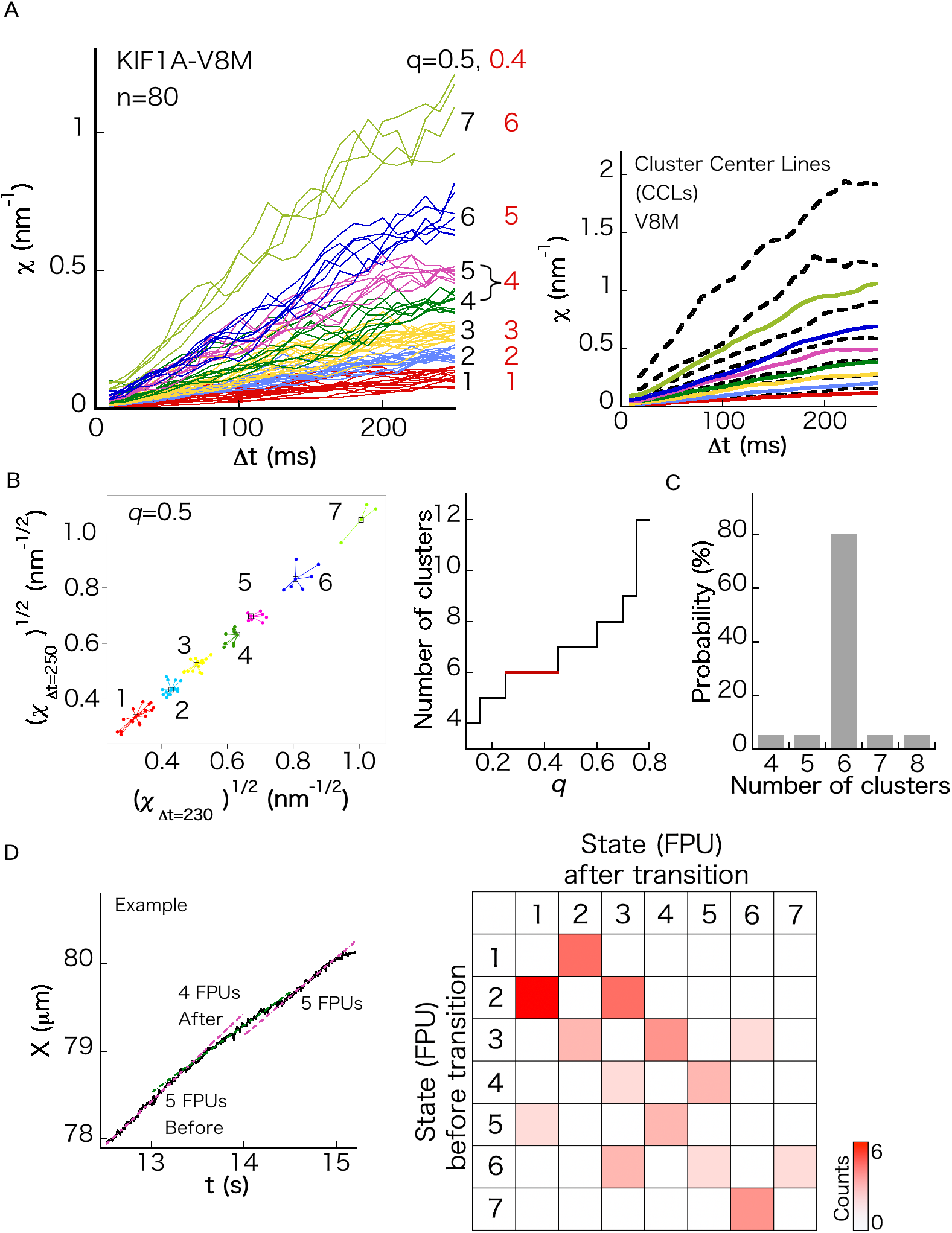
Force index for SVP transport by KIF1A-V8M. (A) Left panel, *χ* as a function of Δ*t* for *n* moving cargos (*n* = 80). Each color represents the cluster classified by the affinity propagation method for *q* = 0.5. The fourth and fifth clusters in case *q* = 0.5 were combined in case *q* = 0.4. Right panel, CCLs calculated as the mean value of each cluster. Black thick lines correspond to KIF1A-WT (Fig. 2A, right panel). (B) Result of affinity propagation analysis (see Materials and Methods). Cluster numbers are plotted as a function of *q*, which is the parameter of the cluster analysis. The most probable cluster number was determined as the cluster number valid for a wide range of *q*. (C) Result of boot-strapping analysis (see Materials and Methods). (D) Left panel, example time course of the center position *X* of a moving SVP cargo, containing velocity change of adjacent CVSs. The straight lines are fitting lines. The number of each FPU at each CVS was determined by calculating *χ*. Right panel, heatmap of transition probability between states (*i.e*., the numbers of FPUs transporting a cargo) of adjacent CVSs.

It has been reported that the V8M mutation relieves the autoinhibitory state of KIF1A so that it does not bind to cargo and consume adenosine triphosphate (ATP) unnecessarily (23). Full-length KIF1A-V8M, which readily takes the active form unlike the full-length KIF1A-WT equipped with an autoinhibitory mechanism, has a microtubule-landing rate of approximately 20-fold that of KIF1A-WT (23). Thus, KIF1A-V8M may attach to SVPs dominantly in the present situation, and a large portion of moving cargos observed in Fig. 4A were considered to be transported by KIF1A-V8M rather than by KIF1A originally derived from mice.

Figures 4B and 4C show the classification results. The fourth and fifth clusters in the case of seven clusters (affinity propagation index q = 0.5) were grouped into the same cluster (Fig. 4A, left panel) in the case of six clusters (q = 0.4). Although six clusters were the most probable based on clustering analyses, seven FPUs transporting SVPs were more plausible according to analysis of the adjacent CVSs (Fig. 4D). As there were some state transitions between the fourth and fifth clusters in the case of q = 0.5 (Fig. 4D, right panel), it was natural not to unite these two clusters as one force unit. Note that back-and-forth transitions (*e.g*., 5 FPUs → 4 FPUs → 5 FPUs) were often observed (Fig. 4D, right panel).

We expected the number of FPUs for KIF1A-V8M to increase because the variant lacks the autoinhibitory mechanism (23), which would lead to an increase in the number of active KIF1A-V8M motors that could attach to SVPs. However, the number of FPUs for KIF1A-V8M was nearly the same as that for KIF1A-WT. This may be because the number of FPUs simultaneously transporting a cargo nearly reached the maximum even for KIF1A-WT in the present experiment, as it was expressed in the cells, which may have resulted in an excess amount of KIF1A in each cell.

### Force index for SVP transport by KIF1A-R350G

For KIF1A-R350G (22), known to cause mutation-associated hereditary spastic paraplegia and to relieve the autoinhibitory mechanism like KIF1A-V8M (21, 23), we did not observe any force impairment. The results are summarized in Fig. S2 in the Supplemental Information. The behavior of *χ* was similar to that of the wild type (Fig. 2A).

### Force index for SVP transport by KIF1A-A255V

KIF1A-A255V (22) is also known to cause mutation-associated hereditary spastic paraplegia and to relieve the autoinhibitory mechanism (23). The results for this variant are summarized in Fig. S3 in the Supplemental Information. As KIF1A-A255V was activated to a lesser extent (half the level of KIF1A-V8M) (23) and *χ* largely reflected the contribution from KIF1A originally derived from mice, the data of *χ* in Fig. S3A were so noisy that they could not be classified. Therefore, to sort at least 1 FPU, *χ*(Δ*t* = 250 ms) < 0.3 nm^−1^ was additionally measured and investigated (Fig. S3A, right panel). For *χ*(Δ*t* = 250 ms) < 0.3 nm^−1^, four clusters were identified through classification (Fig. S3B and S3C). The first and second clusters/third and fourth clusters may be better combined, as we found no transitions between the first and second clusters and third and fourth clusters by adjacent CVS analysis (Fig. S3D). When there are no transitions between these adjacent clusters (as in our observation), it is more natural to consider the number of FPUs to be the same for both clusters. One FPU seemed to contain two clusters, indicating the possible existence of FPUs comprised of different types of dimers (*e.g.*, KIF1A-A255V dimers and dimers of KIF1A-A255V and KIF1A originally derived from mouse).

Figure S3E in the Supplemental Information shows representative kymographs of SVP transport by KIF1A-A255V, KIF1A-WT, KIF1A-V8M, and KIF1A-R350G in the axons of mouse neurons. Frequent anterograde/retrograde reversals of cargos were observed only for KIF1A-A255V. The reason for these reversals has not been elucidated yet, but it should be noted that the first cluster of 1 FPU shown in Fig. S3A, right panel showed the lowest *χ* value, and KIF1A-A255V may lose the tug-of-war with dynein more frequently than other KIF1A variants. Such frequent reversals have not been found in the axons of *C. elegans* neurons (23). This may be because the sizes of SVP cargos are smaller in the axons of *C. elegans* (18), resulting in no significant effect of force impairment under such a low load condition.

### Force index for mitochondrial anterograde transport

The results of recent *in-vitro* single-molecule studies using optical tweezers showed that the force of KIF1A-WT was smaller than that of KIF5 (10, 11). As mitochondria are mainly transported anterogradely by KIF5 (1), we compared *χ* for KIF5-mediated mitochondrial transport (Fig. 5A, left panel) with that for SVP transport by KIF1A-WT. We used the same type of mouse neurons used to observe SVP transport so that the environment of both transports was the same. We selected mitochondria smaller than 1 μm (Fig. 5A, right top panel) to avoid the effect of deformation of large mitochondria when calculating their center positions.

**FIGURE 5.**
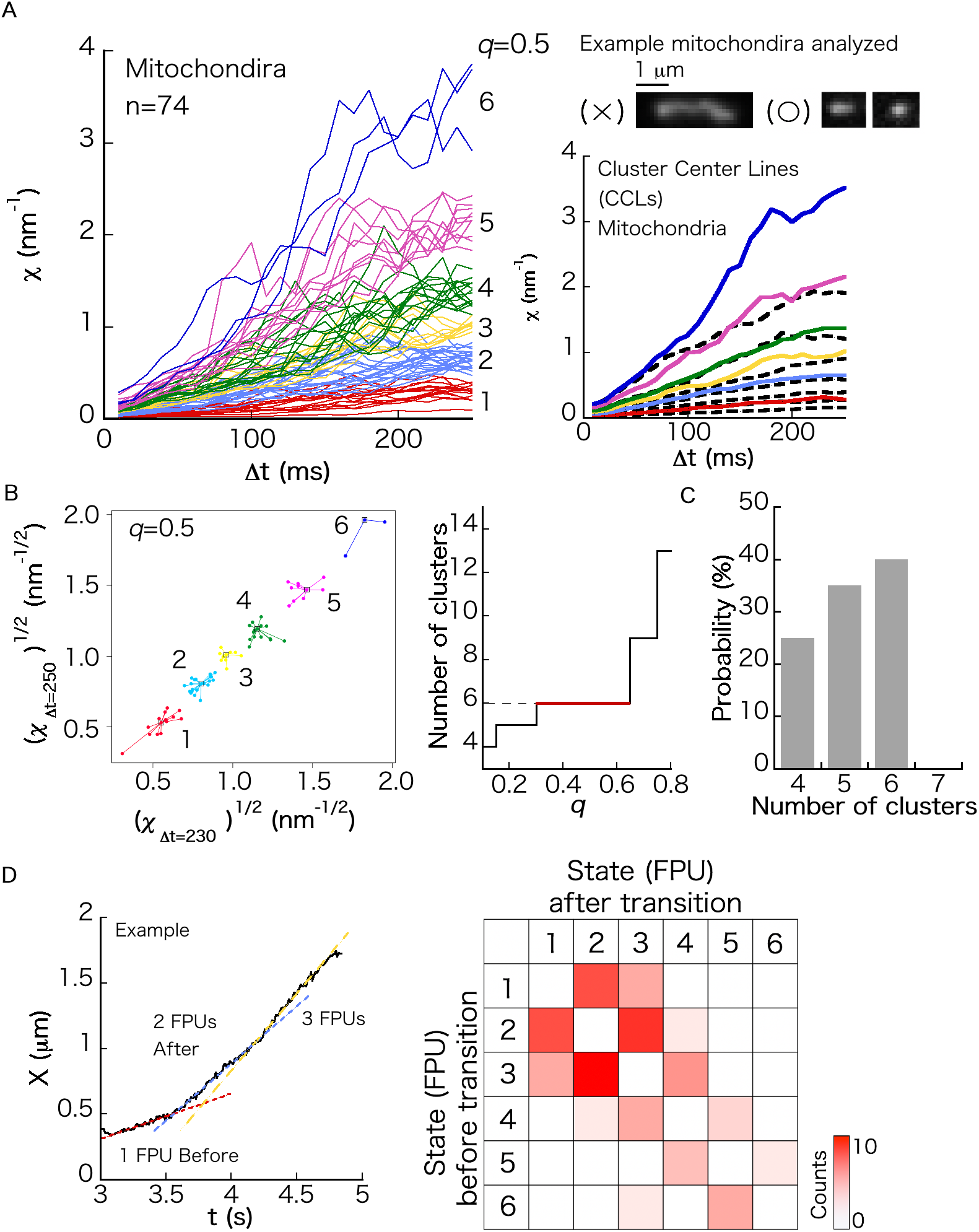
Force index for mitochondria anterograde transport mainly by KIF5. (A) Left panel, *χ* as a function of Δ*t* for *n* moving cargos (*n* = 74). Each color represents the cluster assigned to each FPU. Right panel, CCLs calculated as the mean value of each cluster. Black thick lines correspond to KIF1A-WT (Fig. 2A, right panel). Mitochondria of less than 1 micron were analyzed (micrographs, right top panel). (B) Result of affinity propagation analysis (see Materials and Methods). Cluster numbers are plotted as a function of *q*, which is the parameter of the cluster analysis. The most probable cluster number was determined as the cluster number valid for a wide range of *q*. (C) Result of boot-strapping analysis (see Materials and Methods). (D) Left panel, example time course of the center position *X* of a moving SVP cargo, containing a velocity change of adjacent CVSs. The straight lines are fitting lines. The number of each FPU at each CVS was determined by calculating *χ*. Right panel, heatmap of transition probability between states (*i.e.*, the numbers of FPUs transporting a cargo) of adjacent CVSs.

**FIGURE 6.**
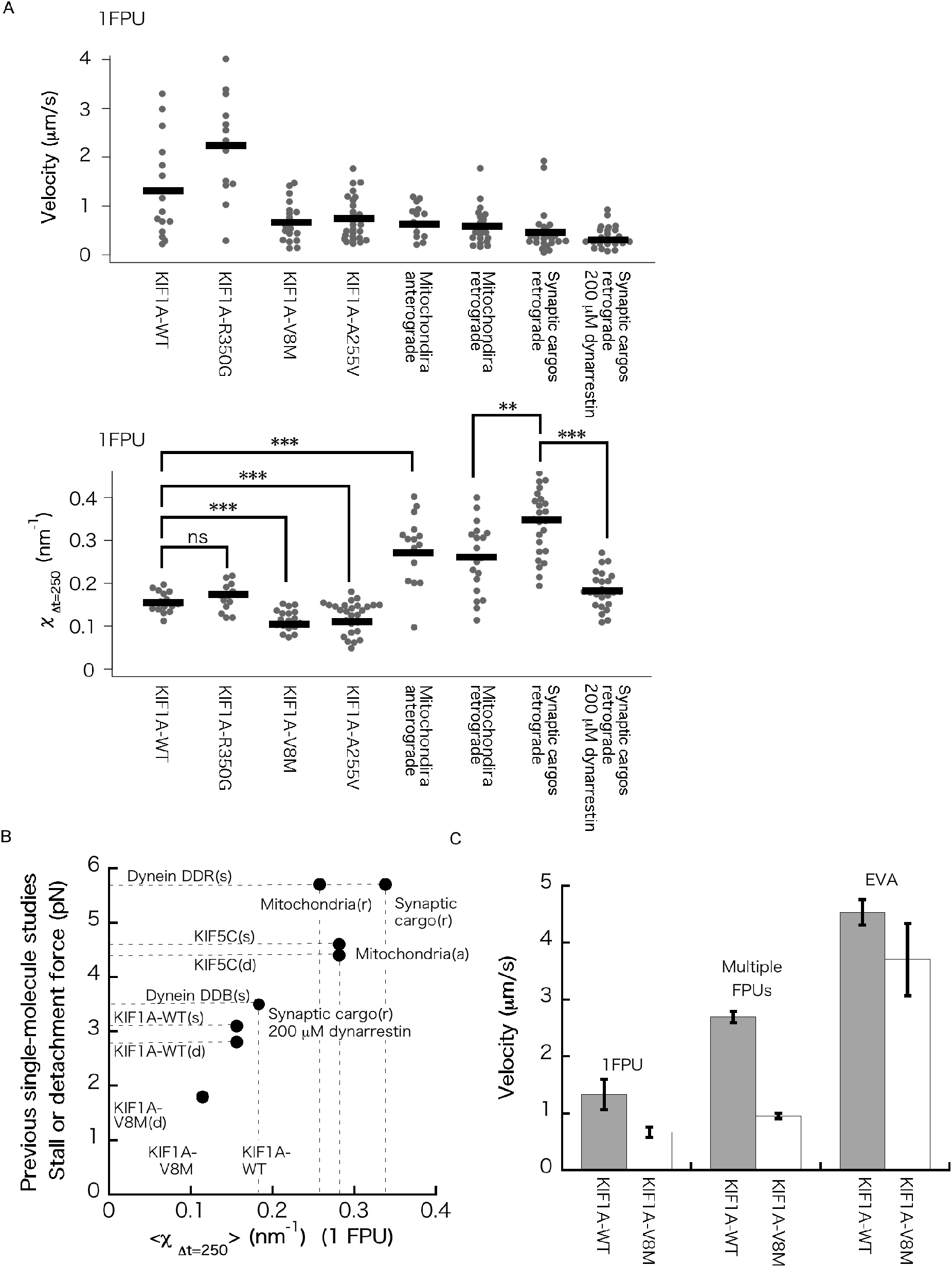
Comparison of force and velocity among different motors. (A) Intracellular velocity (left panel) and force index *χ* (for Δ*t* = 250 ms) in the case of 1 FPU (right panel). \**p* < 0.05, \*\**p* < 0.01, and \*\*\**p* < 0.001. (B) Intracellular force index *χ* in the case of 1 FPU compared with the stall/detachment force measured in single-molecule optical tweezers experiments. 〈*·*〉 represents the mean. The statistical errors of the data are provided in Table S1 in the Supplemental Information. (s) and (d) represent stall force and detachment force, respectively. (a) and (r) represent anterograde and retrograde transport, respectively. KIF1A-WT (s)(d) (full-length motor), KIF1A-V8M (d) (full-length motor) and KIF5C(1–560) (s)(d) have been reported in (11), dynein DDB (one dynein recruited to dynactin) (s) and dynein DDR (two dyneins recruited to dynactin) (s) have been reported in (14). (C) Comparison of the velocity of a moving SVP transported by KIF1A-WT or KIF1A-V8M in case of a mean velocity for 1 FPU, the mean velocity of all data, including multiple FPUs, and the extreme velocity values estimated by EVA (see Materials and Methods, Fig. S7 in the Supplemental Information).

Six clusters were identified for anterograde mitochondrial transport (Fig. 5A, left panel) using the classification (Fig. 5B and 5C). As shown in Fig. 5A, right panel, the *χ* values were larger than those for SVP transport by KIF1A-WT (Fig. 2A), indicating that KIF5 can generate a larger force than KIF1A, as expected from the single-molecule studies (10, 11).

We investigated adjacent CVSs to check whether the clusters in Fig. 5A properly represented the numbers of FPUs. As seen in the representative trajectory (Fig. 5D, left panel), the velocity changes occurring between adjacent numbers of FPUs supported the plausibility of the classification, as it was considered natural for the number to change by ± 1 (Fig. 5D, right panel). Note that a gradual increase (1 FPU→ 2 FPUs→ 3 FPUs) in the number of FPUs was sometimes observed (Fig. 5D, left panel). Transitions between 2 FPUs and 3 FPUs frequently occurred for mitochondrial transport (Fig. 5D, right panel), whereas transitions between 1 FPU and 2 FPUs were dominant in the case of SVPs (Fig. 2D, right panel). Because mitochondria are larger than SVPs, the number of motors attached to mitochondria may be larger.

Mitochondria are important organelles that function as ATP production factories. In neurons, they have to be delivered to areas with high metabolic demand, such as synapses. Abnormalities in mitochondrial transport are strongly associated with diseases, such as Parkinson’s disease and Alzheimer’s disease. The attachment of KIF5 to mitochondria is regulated by mitochondrial adaptors, such as Miro, Milton, and Parkin, which are sensitive to various cellular activities, including an elevated intracellular Ca^2+^ concentration due to neuronal activity (44), and damage caused by reactive oxygen species (45). Therefore, the number of KIF5 motors carrying a mitochondrion can serve as a good biomarker to quantify abnormalities in mitochondria-related cellular phenomena. Future studies should investigate mitochondrial diseases using biomarkers, *i.e.*, the number of KIF5 motors transporting a mitochondrion determined using *χ*.

### Force index for retrograde transport of synaptic cargos and mitochondria

Cytoplasmic dynein is responsible for retrograde cargo transport (3). We investigated synaptic cargo transport by dynein (Fig. S4 in the Supplemental Information). The time course data analyzed here were obtained in our previous study (18). In this reanalysis, *χ* was calculated for Δ*t* = 250 ms (previously, Δ*t* = 150 ms). As *χ* values for higher numbers of FPUs were not classified clearly, we classified *χ*(Δ*t* = 250 ms) < 1.5 nm^−1^, and succeeded in sorting 1 FPU for retrograde synaptic cargo transport (Fig. S4A). The classification (Fig. S4B and S4C) was considered to be appropriate based on the transitions of FPU numbers of adjacent CVSs (Fig. S4D), in which FPU numbers mostly changed by ±1.

Previously, we reported that the value of *χ* for retrograde transport by 1 FPU decreased by half in the presence of the dynein inhibitor, dynarrestin (46). This indicated that the minimal unit to generate force was half of 1 FPU in the absence of the inhibitor (18). A reanalysis of the data in the presence of 200 μM dynarrestin is shown in Fig. S5 in the Supplemental Information. The value of *χ* for the sorted 1 retrograde FPU in the presence of 200 μM dynarrestin was half of that in the absence of the inhibitor (Fig. S4A).

Retrograde transport of mitochondria was also investigated (Fig. S6 in the Supplemental Information). We found that the value of 1 retrograde FPU for mitochondria did not largely differ from that for synaptic cargos. Although the cargo types were different, the force generation of 1 retrograde FPU generated by cytoplasmic dynein was similar.

### Comparison of *χ* among intracellular cargo transport by different microtubule motors

Finally, we summarized the results of the measurements of *χ*. Figure 6A shows comparisons of the velocity and force index *χ*(Δ*t* = 250ms) in the case of the sorted 1 FPU for different types of cargo transport by different microtubule motors. While velocity (Fig. 6, left panel) does not accurately represent the motor’s ability because it largely depends on cargo size (see Fig. S1B for the various sizes), the force index *χ* of 1 FPU, which depends less on cargo size, was considered to represent the ability of the motors.

Although *χ* is an index of the drag force acting on a moving cargo, its value is close to the stall/detachment force when the load acting on the motor is sufficiently large in the intracellular environment. We compared the values of *χ* in the case of 1 FPU with the stall/detachment force values of full-length motors measured in *in-vitro* single-molecule experiments using optical tweezers (11, 14) (Fig. 6B). In the case of dynein, we compared 1 retrograde FPU in the absence or presence of dynarrestin with two dynein dimers/one dynein dimer attached to dynactin (14). There seemed to be a high correlation between the *χ* value of 1 FPU and the stall/detachment force of microtubule motors measured in the single-molecule experiments.

### EVA of transport velocity

Determinations of the velocity of KIF1A-V8M have led to controversial results (11, 23). In our experiment, the mean velocity of moving SVPs transported by 1 FPU of KIF1A-V8M was 50 lower than that of SVPs transported KIF1A-WT (Fig. 6C). A similar behavior was reported by Budaitis *et al*. (11), who observed a 60% decrease in the velocity of KIF1A-V8M in single-molecule TIRF assays. When we calculated the mean velocities using all data combined, including transport by multiple FPUs, the velocity values for both motors slightly increased (Fig. 6C), suggesting that the load to each motor was decreased through multiple-motor cooperation. For intracellular cargo transport, the velocity of the motor is load-dependent.

To determine the velocity under low load conditions near the maximum velocity of the motors, we subjected the velocity data to EVA (24) (Fig. 6C). We expected to obtain a velocity close to the value under the limiting condition in which the cargo size is the smallest. The distributions of the extreme velocity values are provided in Fig. S7 in the Supplemental Information. The EVA results in Fig. 6C, indicate that the velocity of KIF1A-V8M was close to that of KIF1A-WT. Because the force generation of KIF1A-V8M is impaired (11), the motor movement was sensitive to the load. Chiba *et al.* (23) reported that the mean velocity values of moving SVPs in *C. elegans* axons were similar for UNC104-WT and UNC104-V6M where UNC-104 is a homolog of KIF1A. This may be because the SVPs in *C. elegans* axons are substantially smaller than those in mouse neuronal axons (18), and the effect of force impairment was not significant under the low-load condition realized in *C. elegans* axons.

Lastly, we investigated the effect of the presence of KIF1A originally derived from mice on EVA (Fig. S7 in the Supplemental Information). Judging from the return level plots of KIF1A-V8M and KIF1A-A255V, the data for KIF1A-A255V alone were considered to actually contain data of KIF1A originally derived from mice, as the convergence value changed from the middle to the convergence value of KIF1A-WT. Note that the return level plot of EVA is roughly similar to the cumulative distribution of extreme velocity values, and the convergence value is the extreme value. As KIF1A-V8M showed a single convergence value in the return level plot, the measured velocities for KIF1A-V8M did not include much the velocity data of KIF1A originating from mice.

## Conclusion

We measured the force index *χ*, and compared *χ* values in the case of 1 FPU with motor stall/detachment forces obtained by *in-vitro* single-molecule experiments (11, 14). We found a strong correlation between them (Fig. 6B), indicating the validity of the non-invasive force measurement method based on the force index *χ*. To our knowledge, this was the first study to investigate the pathogenic variants KIF1A-V8M (21), KIF1A-R350G and KIF1A-A255V (22), which cause hereditary spastic paraplegia, in the axons of mammalian neurons (Fig. 6A and 6C). Force and velocity impairment of the V8M and A255V variants were observed in mammalian neurons, unlike in *C. elegans* neurons (23), possibly because the high load condition that seriously affects motor force impairment is realized only in axonal transport in mammalian neurons (Fig. 6A, 6C and S3E). Finally, we note that this study was the first to apply EVA to axonal transport to extract near-maximum velocity values of the motors without cargos. We hope the time course analyses developed in the present paper are helpful for future studies on neuronal diseases, including KAND.

## Author Contributions

K. H., S. M., and Y. I. performed the experiments. K. H. and T. N. analyzed the data. K. H. wrote the paper.

## Acknowledgements

We acknowledge Dr. S. Niwa for providing the plasmids of the KIF1A variants and fruitful discussions on SVP transport, including an initial idea on the application of EVA. We also acknowledge M.G. Miyamoto and K. Nagino for experimental support. We would like to thank Edita<u>g</u>e (www.editage.com) for English language editing. This work was supported by JST PRESTO (grant No. JPMJPR1877), AMED PRIME (grant No. JP18gm5810009), JSPS KAKENHI (grant No. 17H03659), and FRIS Creative Interdisciplinary Research Program Tohoku University to K. H.

## Data Availability

Data supporting the findings of this study are available within the article and its Supplemental Information files and from the corresponding author upon reasonable request.

## Supplemental Information

**FIGURE S1.**
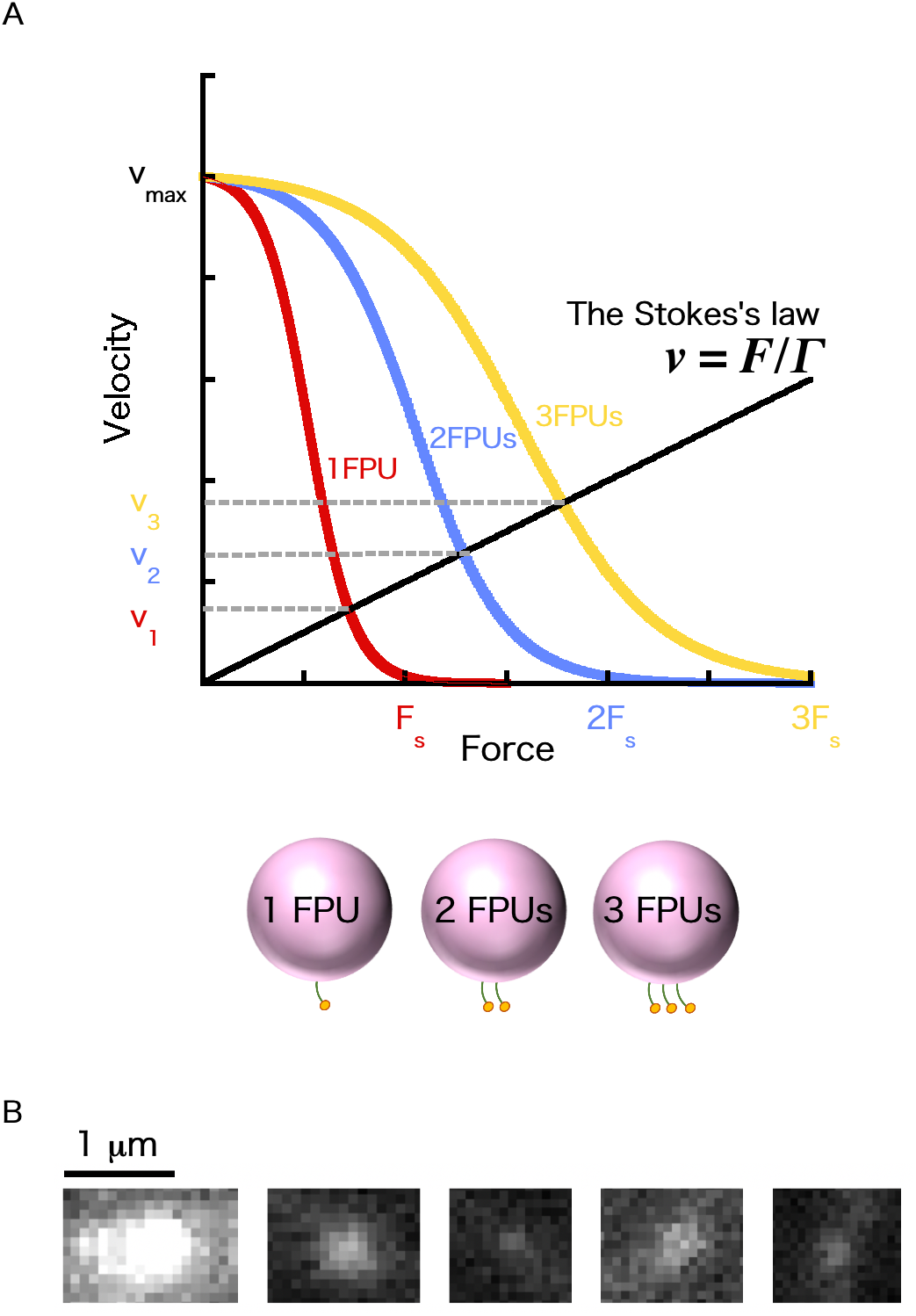
Relationships among motor velocity, force, and cargo size. (A) Schematic representation of force-velocity relationships of kinesin in the cases of 1 FPU (1 dimer), 2 FPUs (2 dimers), and 3 FPUs (3 dimers), based on a previous study (1). The straight line represents Stokes’ law, where *χ* is the velocity of a moving cargo, *F* is the force exerted by the motors, and Γ is the friction coefficient of the cargo. *v*_1_, *v*_2_, *v*_3_ represent the velocities of the same cargo in the cases of 1 FPU (1 dimer), 2 FPUs (2 dimers) and 3 FPUs (3 dimers), determined by the intersections of the force-velocity relationship and Stokes’ law. *F*_s_ is the stall force of the motor, and *V*_max_ is the maximum velocity. The slope (1/Γ) of the line is different for each cargo because their sizes (∝ Γ) differ. (B) Representative fluorescence micrographs of synaptic vesicle precursor (SVP) cargos of various sizes observed in the axons of mouse neurons.

**FIGURE S2.**
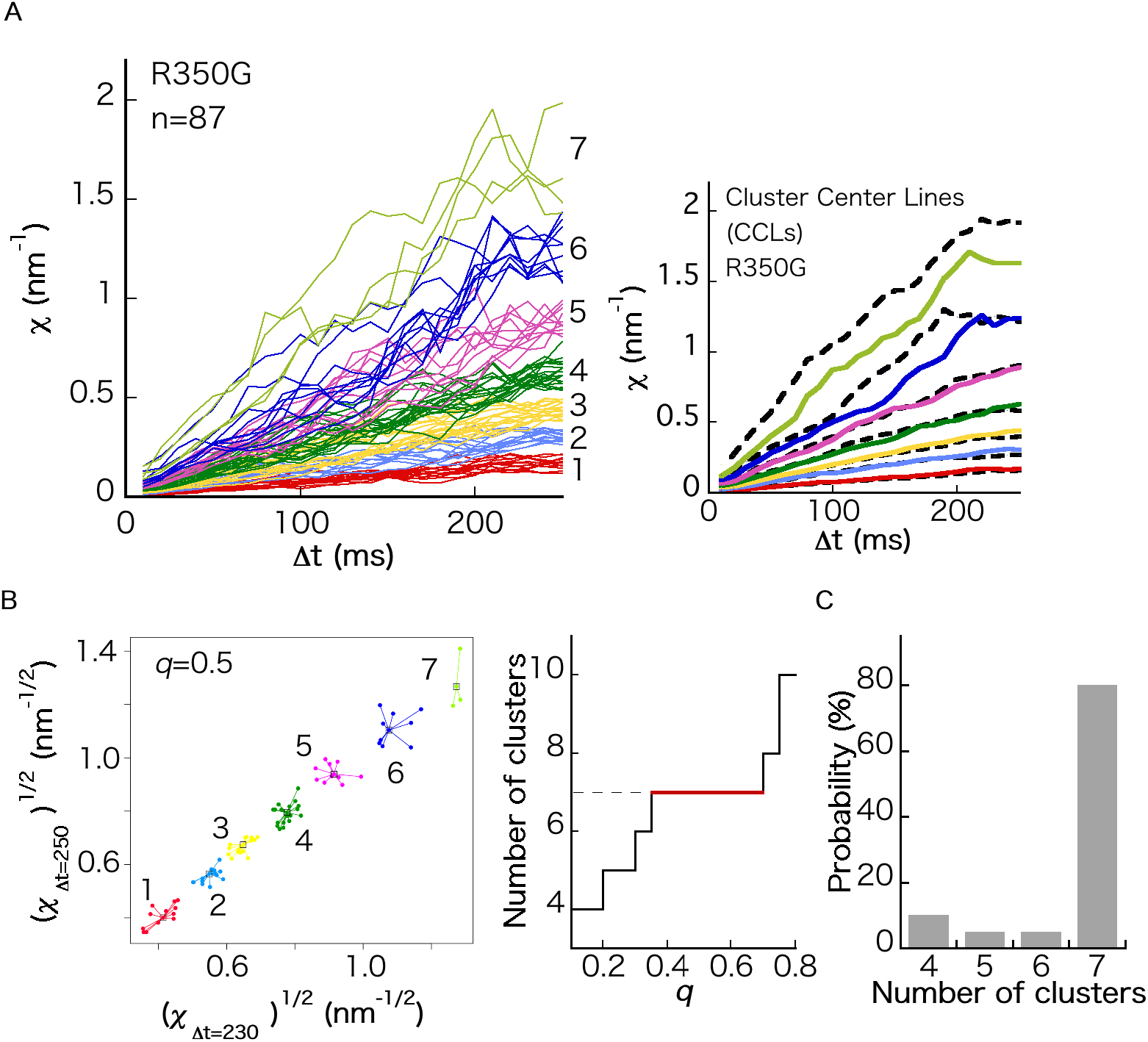
Force index for SVP transport by KIF1A-R350G. (A) Left panel, *χ* as a function of Δ*t* for *n* moving cargos (*n* = 87). Each color represents the cluster assigned to each FPU. Right panel, cluster center lines (CCLs) calculated as the mean value of each cluster. The black thick lines correspond to KIF1A-WT (Fig. 2A in the main text, right panel). (B) Result of affinity propagation analysis (see Materials and Methods). Cluster numbers are plotted as a function of *q*, which is the parameter of the cluster analysis. The most probable cluster number was determined as the cluster number valid for a wide range of *q*. (C) Result of boot-strapping analysis (see Materials and Methods).

**FIGURE S3.**
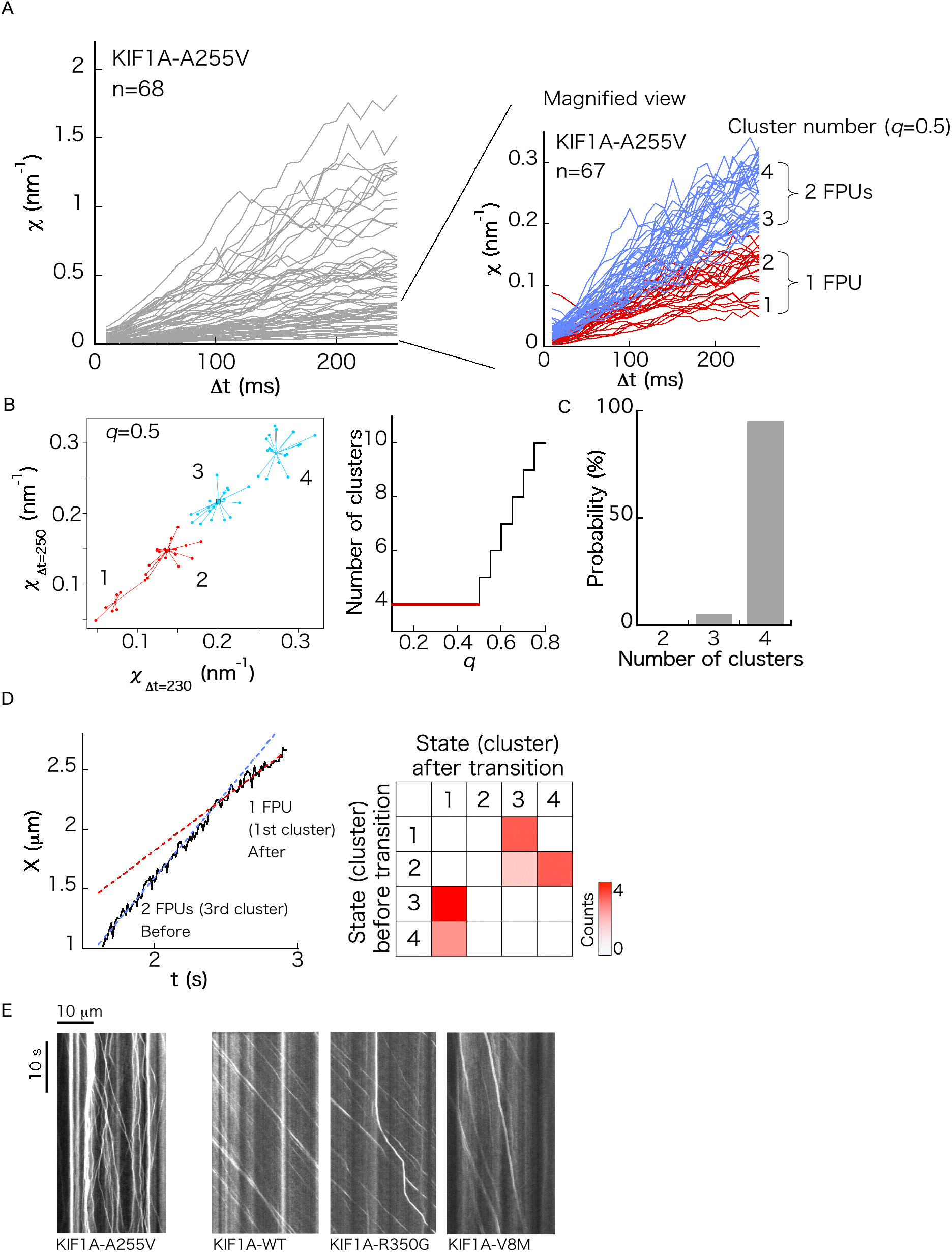
Force index for SVP transport by KIF1A-A255V. (A) Left panel, *χ* as a function of Δ*t* for *n* moving cargos (*n* = 68). Each color represents the cluster assigned to each FPU. To sort 1 FPU cluster, *χ* < 0.3 nm^−1^ was additionally measured and investigated (right panel, *n* = 67). Each color represents the number of FPUs, each of which contains two clusters. (B) Result of the affinity propagation analysis (see Materials and Methods). Cluster numbers are plotted as a function of *q*, which is the parameter of the cluster analysis. The most probable cluster number was determined as the cluster number valid for the wide range of *q*. (C) Result of the boot-strapping analysis (see Materials and Methods). (D) Left panel, representative time course of the center position *X* of a moving SVP cargo, containing a velocity change of adjacent CVSs. The straight lines are fitting lines. The number of each FPU at each CVS was determined by calculating *χ*. Right panel, heatmap of transition probability between states (*i.e.,* the numbers of FPUs transporting a cargo) of adjacent CVSs. Because we observed no 1st↔cluster 2nd↔cluster and 3rd cluster↔4th cluster transitions, 1st and 2nd clusters and 3rd and 4th clusters were combined as the same clusters. (E) Representative kymographs of SVP transport by KIF1A-A255V, KIF1A-WT, KIF1A-V8M, and KIF1A-R350G in the axons of mouse neurons.

**FIGURE S4.**
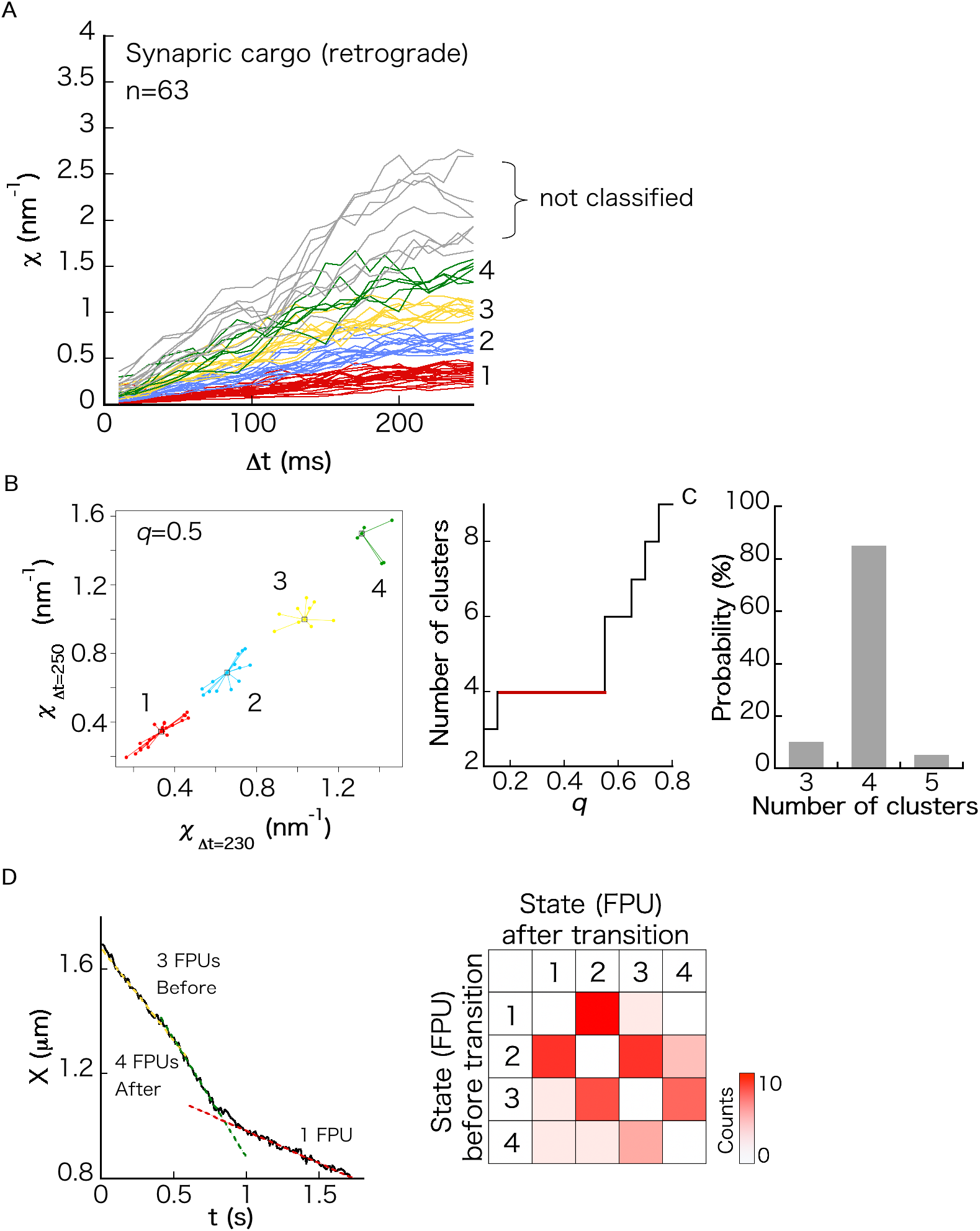
Force index *χ* for retrograde transport of synaptic cargos. (A) Left panel, *χ* as a function of Δ*t* for *n* moving cargos (*n* = 63). Each color represents the cluster assigned to each FPU. The time course data were obtained in our previous study (2). In this reanalysis, *χ* was calculated until Δ*t* = 250 ms (previously, it was calculated until Δ*t* = 150 ms). Because *χ* values for higher numbers of FPUs were not observed clearly, we classified *χ*(Δ*t* = 250 ms) < 1.5 nm^−1^. (B) Result of affinity propagation analysis (see Materials and Methods). Cluster numbers are plotted as a function of *q*, which is the parameter of the cluster analysis. The most probable cluster number was determined as the cluster number valid for the wide range of *q*. (C) Result of boot-strapping analysis (see Materials and Methods). (D) Left panel, representative time course of the center position *X* of a moving SVP cargo, containing velocity change of adjacent CVSs. The straight lines are fitting lines. The number of each FPU at each CVS was determined by calculating *χ*. Right panel, heatmap of transition probability between states (*i.e*., the numbers of FPUs transporting a cargo) of adjacent CVSs.

**FIGURE S5.**
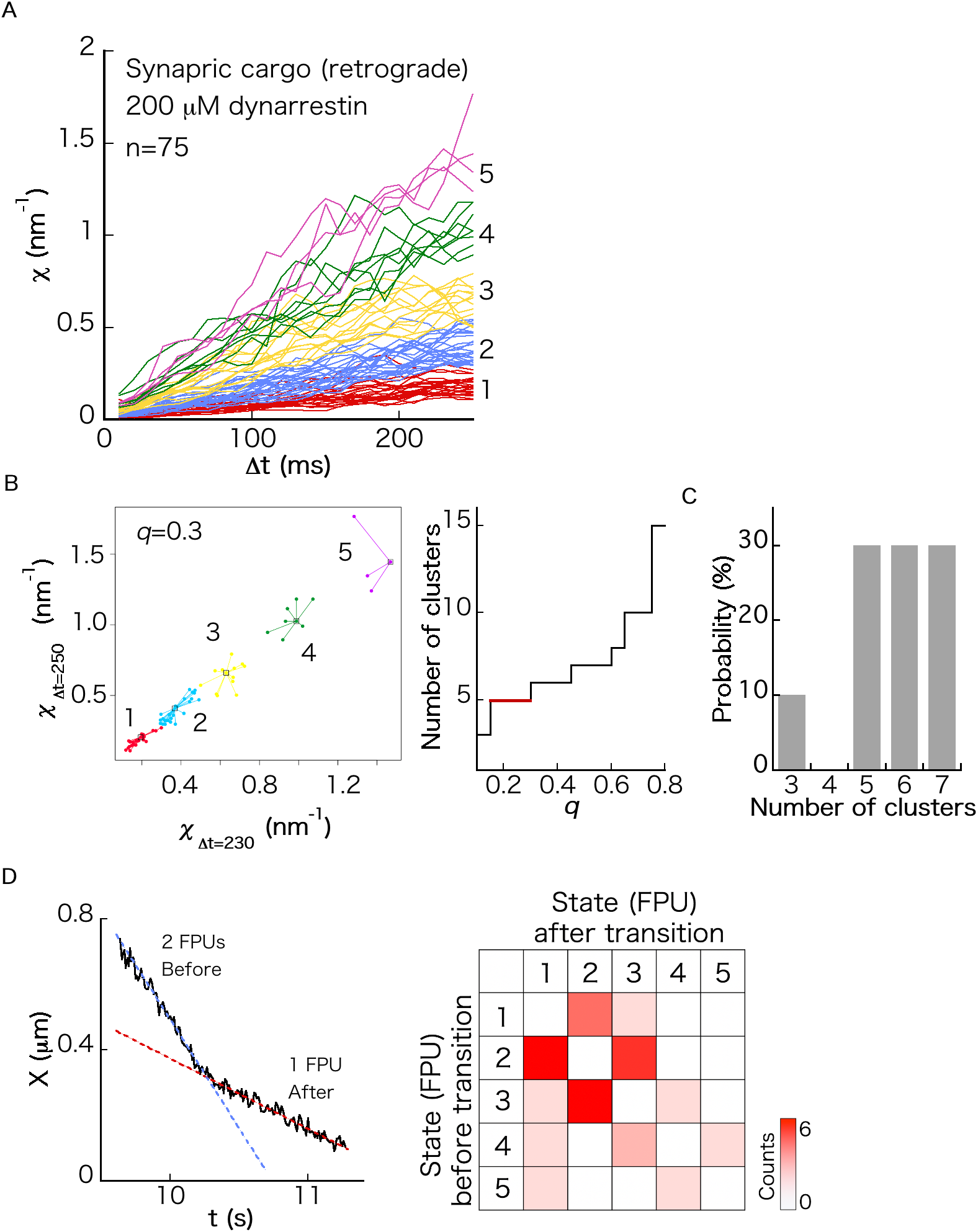
Force index for retrograde transport of synaptic cargos in the presence of 200 μM dynarrestin. (A) Left panel, *χ* as a function of Δ*t* for *n* moving cargos (*n* = 75). Each color represents the cluster assigned to each FPU. The time course data were obtained in our previous study (2). In this reanalysis, *χ* was calculated until Δ*t* = 250 ms (previously, it was calculated until Δ*t* = 150 ms). (B) Result of affinity propagation analysis (see Materials and Methods). Cluster numbers are plotted as a function of *q*, which is the parameter of the cluster analysis. The most probable cluster number was determined as the cluster number valid for the wide range of *q*. (C) Result of boot-strapping analysis (see Materials and Methods). (D) Left panel, representative time course of the center position *X* of a moving SVP cargo, containing a velocity change of adjacent CVSs. The straight lines are fitting lines. The number of each FPU at each CVS was determined by calculating *χ*. Right panel, heatmap of transition probability between states (*i.e*., the numbers of FPUs transporting a cargo) of adjacent CVSs.

**FIGURE S6.**
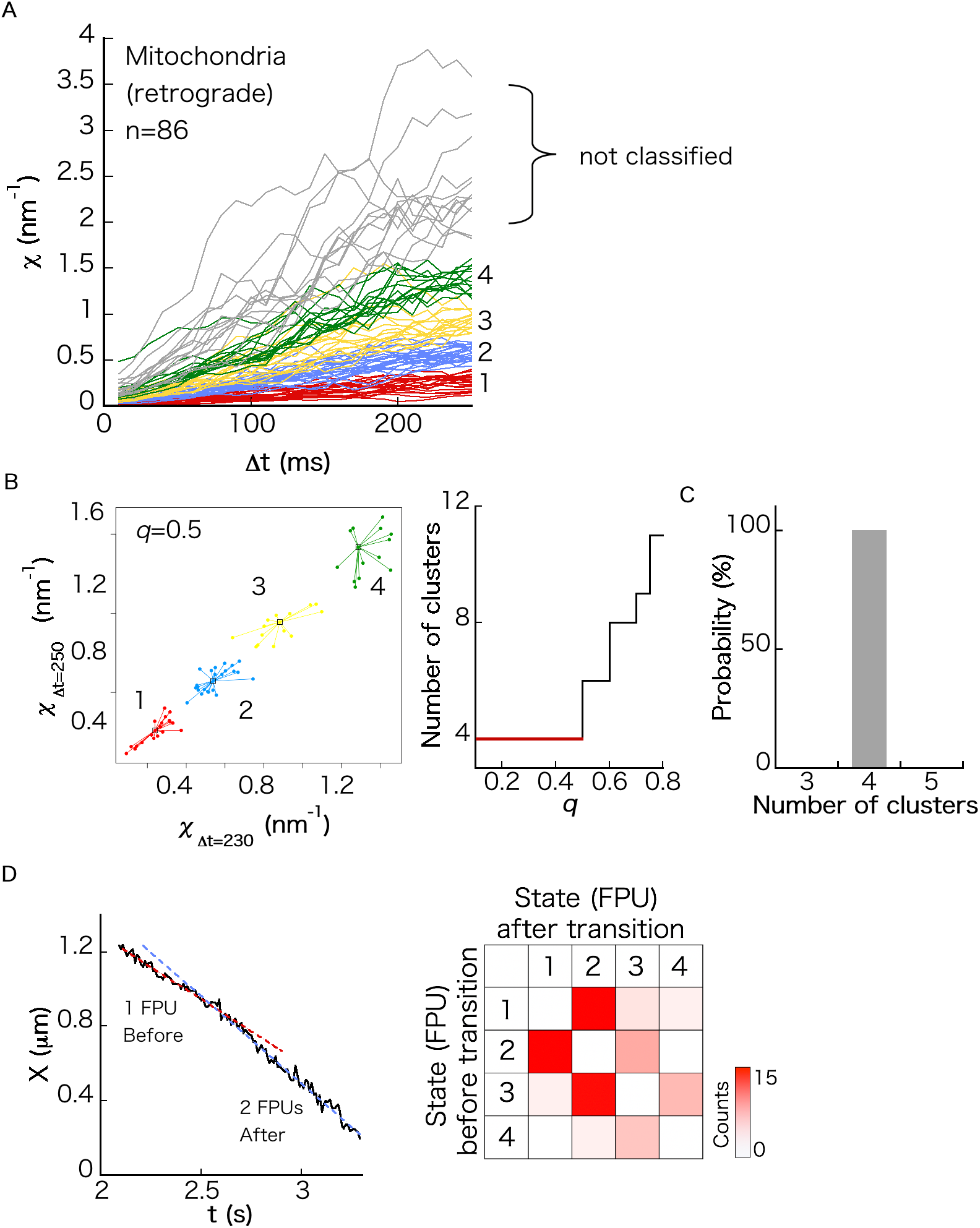
Force index for retrograde transport of mitochondria. (A) Left panel, *χ* as a function of Δ*t* for *n* moving cargos (*n* = 86). Each color represents the cluster assigned to each FPU. Right panel, CCLs calculated as the mean value of each cluster. As *χ* values for higher numbers of FPUs were not observed clearly, we classified *χ*(Δ*t* = 250 ms) < 1.5 nm^−1^. (B) Result of affinity propagation analysis (see Materials and Methods). Cluster numbers are plotted as a function of *q*, which is the parameter of the cluster analysis. The most probable cluster number was determined as the cluster number valid for a wide range of *q*. (C) Result of boot-strapping analysis (see Materials and Methods). (D) Left panel, representative time course of the center position *X* of a moving SVP cargo, containing a velocity change of adjacent CVSs. The straight lines are fitting lines. The number of each FPUs at each CVS was decided by calculating *χ*. Right panel, heatmap of transition probability between states (*i.e*., the numbers of FPUs transporting a cargo) of adjacent CVSs.

**FIGURE S7.**
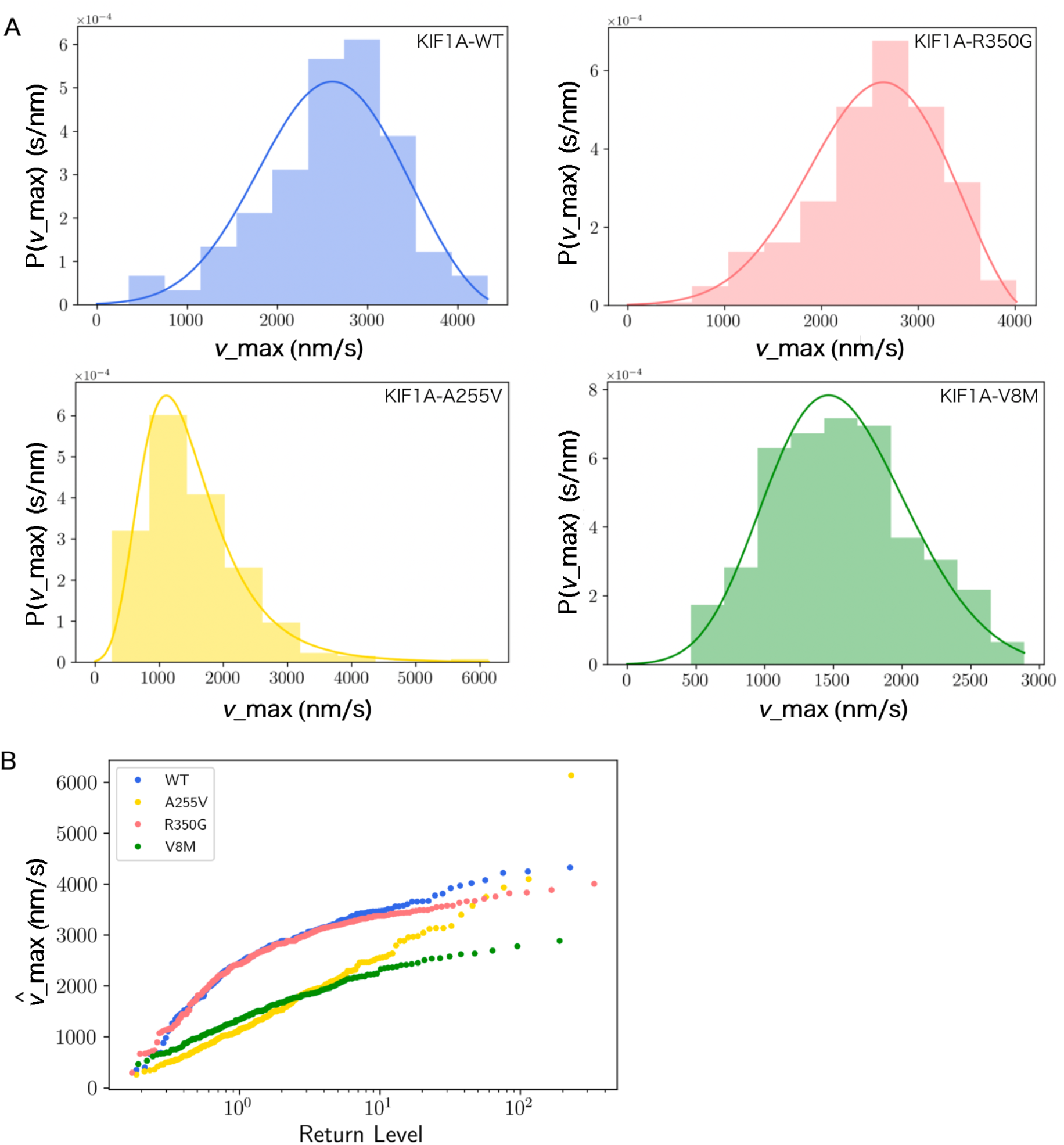
Extreme value analysis for SVP transport. (A) Probability distributions *g*(*v*_max_) of *v*_max_ of SVP transport velocity. Fitted lines represent the derivatives of Eq. 4 of the main text (Materials and Methods). (B) Return level plot defined as 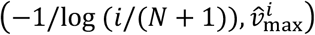, where 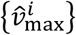 is the rearranged data of 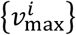 so that 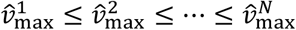.

**Table S1.** Stall and detachment forces reported in previous single-molecule studies.

| Motor | Force | Reference |
| --- | --- | --- |
| KIF1A-WT(FL) (s) | $2.98 \pm 0.11$ pN | (3) |
| KIF1A-WT(FL) (d) | 2.8 (2.4, 3.3) pN | (3) |
| KIF1A-V8M(FL) (d) | 1.8 (1.5, 2.1) pN | (3) |
| KIF5C(1-560) (s) | $4.6 \pm 0.04$ pN | (3) |
| KIF5C(1-560) (d) | 4.4 (3.8, 4.9) pN | (3) |
| dynein DDR (s) | $5.7 \pm 0.2$ pN | (4) |
| dynein DDB (s) | $3.6 \pm 0.1$ pN | (4) |

FL: full length, ( ): quartiles, (s): stall force, (d):detachment force, mean ± SEM.

**Force index $\langle \chi_{\Delta t=250} \rangle$ for intracellular cargo transport by 1 FPU**
| Transport | $\langle \chi_{\Delta t=250} \rangle$ | Reference |
| --- | --- | --- |
| SVP transport by KIF1A-WT | $0.156 \pm 0.00063$ nm<sup>-1</sup> | This study |
| SVP transport by KIF1A-V8M | $0.114 \pm 0.00063$ nm<sup>-1</sup> | This study |
| Mitochondrial transport (a) | $0.282 \pm 0.00077$ nm<sup>-1</sup> | This study |
| Mitochondrial transport (r) | $0.258 \pm 0.00078$ nm<sup>-1</sup> | This study |
| Synaptic cargo transport (r) | $0.338 \pm 0.00074$ nm<sup>-1</sup> | (2) |
| Synaptic cargo transport (r)<br>200 μM dynarrestin | $0.183 \pm 0.00042$ nm<sup>-1</sup> | (2) |

(a): anterograde, (r):retrograde, mean ± SEM

